# 3D epithelial cell topology tunes signalling range to promote precise patterning

**DOI:** 10.1101/2025.08.08.668674

**Authors:** Giulia Paci, Francisco Berkemeier, Buzz Baum, Karen M. Page, Yanlan Mao

## Abstract

Cell behaviours in multicellular organisms are coordinated via both diffusible molecules and by signals based on direct cell-cell contacts. The mode of cell communication used influences the signalling range. In many developing epithelia, contact-based Notch-Delta lateral inhibition signalling is used to pattern cell fates. While previous work revealed that cells can use protrusions to extend the range of Notch-Delta signalling to alter these patterns, this is not a general feature of epithelia. In addition, it is not known how the complex three-dimensional (3D) shapes of epithelial cells influence cell communication. In exploring this question, we show that epithelial cells at the *Drosophila* wing margin, which lack basal protrusions, contact different neighbours at different heights along their apico-basal axis, effectively increasing the number of neighbours each cell touches. To quantitatively assess this behaviour, we develop a novel mathematical modelling framework (Multi-layer Signalling Model) to simulate Notch-Delta signalling over data-derived 3D cell topologies. The model predicts that lateral cell surface signalling is essential to tune the spacing between SOPs. In agreement, we show that perturbing cortical stiffness and cell tortuosity *in vivo* modifies SOP spacing. These results emphasise the importance of 3D cell geometry and topology in fine-tuning signalling range.

## Introduction

Cell-cell signalling is essential for the coordination of cellular behaviours within living organisms, including cell growth, differentiation, immune response, tissue formation and homeostasis. Signalling can occur through diffusible chemicals that are secreted locally or into the circulatory system, and/or via direct cell-cell contacts. In epithelia, cells can be densely packed into elongated and tortuous shapes such as scutoids (1) and punakoids (2), that reflect changes in cell contacts along the apico-basal axis of cells. Although multiple signalling molecules and their receptors can be found on the apical and baso-lateral surfaces of cells (3–5), the direct impact of 3D cell-cell contacts (topology) on cell signalling range is currently unclear.

Fine-tuning cell signalling range is essential for the establishment of precise patterns of cell fates, such as limb digits, hair follicles and sensory cells in the inner ear. These precise structures can be achieved through reaction-diffusion processes (6), morphogen gradients (7) or via direct cell-cell interactions - as is the case for Notch-Delta lateral inhibition (8). In lateral inhibition, an initially mostly homogenous cell population has the potential to acquire either a “signal-sending” or “signal-receiving” fate. Through a self-organisation process, a subset of cells is selected to acquire a “signal-sending” fate and subsequently inhibit this fate from its neighbours (Figure 1a). In recent years, several mechanisms by which cells can extend their range of inhibition have been identified (9), including filopodia (10–13) and dynamic cell rearrangements (14) which contribute to sparser patterning (Figure 1b).

**Figure 1:**
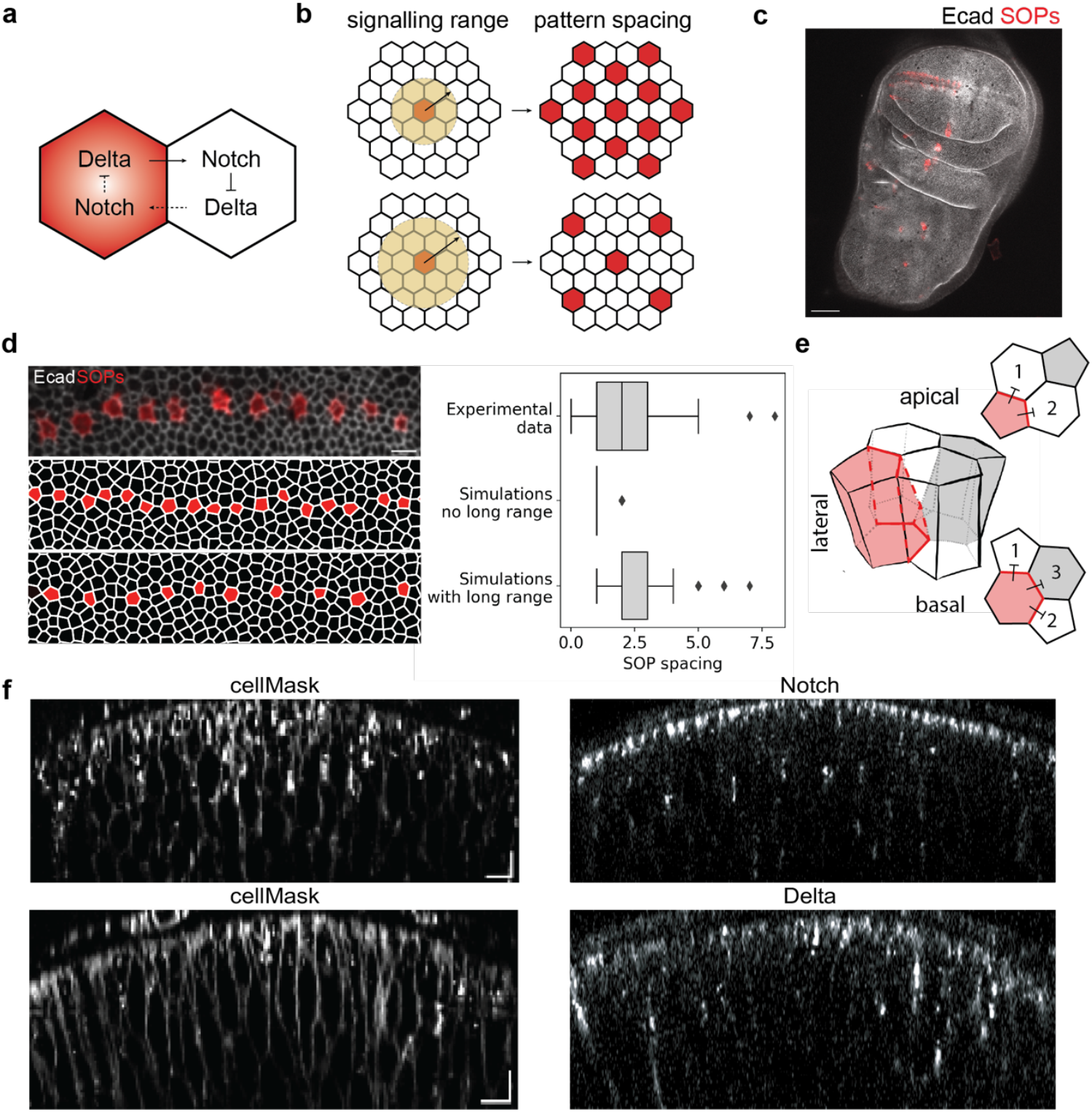
Signalling beyond the first apical neighbours is required to achieve the experimentally observed SOP pattern. **a**, Feedback loop underlying lateral inhibition in Notch-Delta signalling: Delta ligands on the red cell activate Notch in its neighbour, leading to Delta repression. This in turn decreases Notch activation in the red cell leading to a high level of Delta. **b**, For patterning based on lateral inhibition, an extended signalling range gives rise to a sparse pattern as the cells can inhibit neighbours further away in the tissue. **c**, Overview of SOPs in a late third instar wing disc, including the two rows of SOPs at the anterior wing margin, and SOPs in the notum. Scale bar: 50 μm. **d**, Left: SOP pattern in late 3^rd^ instar wing discs (top, scale bar: 5 μm) compared to simulated tissues without (middle) and with (bottom) long-range signalling. Right: quantification of SOP spacing (number of cells between two SOPs) in experiments and simulations. **e**, Simplified schematic of neighbour changes in 3D and their impact on signalling. The four cells undergo an apico-basal intercalation so that the red cell contacts cells 1 and 2 on the apical surface and cells 1,2 and 3 on the basal surface. The apical and basal planes are shown as representative, but similar intercalations will involve larger portions of the lateral surfaces. **f**, Membrane staining (cellMask) and localisation of Delta and Notch in third instar wing discs, side views, scale bar: 5 μm.

Here we uncover a fundamental new mechanism of Notch-Delta signalling. Through a computational model of Notch-Delta signalling we show that cellular apical contacts alone cannot explain the experimentally observed pattern of sensory organ precursors (SOPs) at the *Drosophila* wing margin. We established an imaging pipeline combining two-photon imaging and machine learning assisted segmentation to reconstruct the precise 3D cell shapes in live tissues. By mapping the cell-cell contacts we find that wing disc cells undergo multiple neighbour exchanges along their apico-basal axis. Incorporating the 3D cell topology into a new multi-layer signalling model, we show how these extra contacts on lateral cell surfaces can extend Notch-Delta signalling range to give rise to the precise bristle spacing observed *in vivo*. Finally, we find that directly perturbing the cell shapes to reduce lateral contacts reduces the SOP spacing as predicted. Altogether, these results emphasise the need to consider 3D cell shapes in the wider context of cell-cell communication and prompt us to revisit how signals are relayed within complex tissues.

## Results

### Signalling beyond first apical neighbours is required to achieve the correct patterning of SOPs

We quantified the pattern of SOPs at the wing margin at the time of specification by imaging late third instar wing discs and measuring the distance between successive SOPs (Figures 1c,d). The pattern of SOPs is sparse, with an average spacing of 2-3 epithelial cells between each SOP cell (Figure 1d, top). This contrasts with the salt-and-pepper pattern that would be expected by apical lateral inhibition dynamics alone, where each cell can only inhibit its direct apical neighbours (9).

To quantitatively understand the requirements for sparse patterning, we developed a mathematical modelling framework to simulate the emergence of the SOP pattern through lateral inhibition (Figure 1d). For each cell, we assume juxtacrine signalling with all neighbouring cells, as proposed by Collier et al. (15). In this model, Notch activity is described as an increasing function of the Delta activity in neighbouring cells, while Delta activity is inhibited by Notch within the same cell, creating a negative feedback loop (see (16) and Supplementary Information). Both processes are modulated by a decay term to ensure dynamic stability.

As expected, when cells are allowed to signal only to their first neighbours, a tightly packed salt-and-pepper pattern emerges, with SOPs separated by a single non-SOP cell (Figure 1d, middle). We then extended the model to incorporate long-range signalling and found that including second to third neighbour signalling (signalling range equal to twice the cell diameter) could recapitulate the experimentally observed pattern (Figure 1d, bottom and Extended Data Figure 1). These results indicate that the pattern of wing margin SOPs cannot be explained by conventional lateral inhibition through 2D neighbour contacts alone. Instead, some form of long-range signalling is needed.

Wing disc cells have no visible basolateral filopodia (10) to extend signalling range. Instead, by having an apico-basally elongated shape and extended lateral surface (Figure 1e), we reasoned that signalling range could be increased laterally as cells exchange and contact new neighbours along their apico-basal axis. This effectively increases the number of neighbours each cell directly contacts and the effective signalling range (Figure 1e). However, lateral signalling requires the presence of Notch and Delta along the baso-lateral surface of cells, which has been observed in other systems (10, 17, 18). To check whether lateral contacts could be signalling-competent, we imaged the localisation of Delta and Notch in third instar discs using knock-in lines. In addition to the expected strong apical localisation of Notch, we found that both Notch and Delta are present on lateral cell membranes (Figure 1f).

### Pseudo-stratified wing disc cells undergo several intercalations along the apico-basal axis

To understand how the topology of cell-cell contacts (and therefore inhibition) changes when considering the 3D height of the cells compared to just 2D apical contacts, we developed a new pipeline to image and segment individual cells within live wing discs (Figures 2a,b); see Methods and (19). We found that cells undergo extensive neighbour exchanges (intercalations) along the apico-basal axis, with apical neighbours moving away from each other in z and vice versa (examples of both types in Figure 2c). These 3D intercalations amount to an overall two-fold increase of unique cell contacts when compared to the number of neighbours defined by apical contacts alone (Figure 2d). These intercalations enable cells to effectively extend their neighbourhoods and reach beyond the first surrounding apical neighbours (Figure 2e), with the potential to extend their signalling range. Nearly all cells undergo 3D intercalations, with a slight anti-correlation between the number of starting apical neighbours and the additional number of neighbours gained in z (Extended Data Figure 2a,b). Since wing disc cells are very apico-basally elongated, their lateral surface area is much greater than apical (Extended Data Figure 2c), therefore the total contribution to signalling from this area may be significant despite the reduced concentration of Notch/Delta.

**Figure 2:**
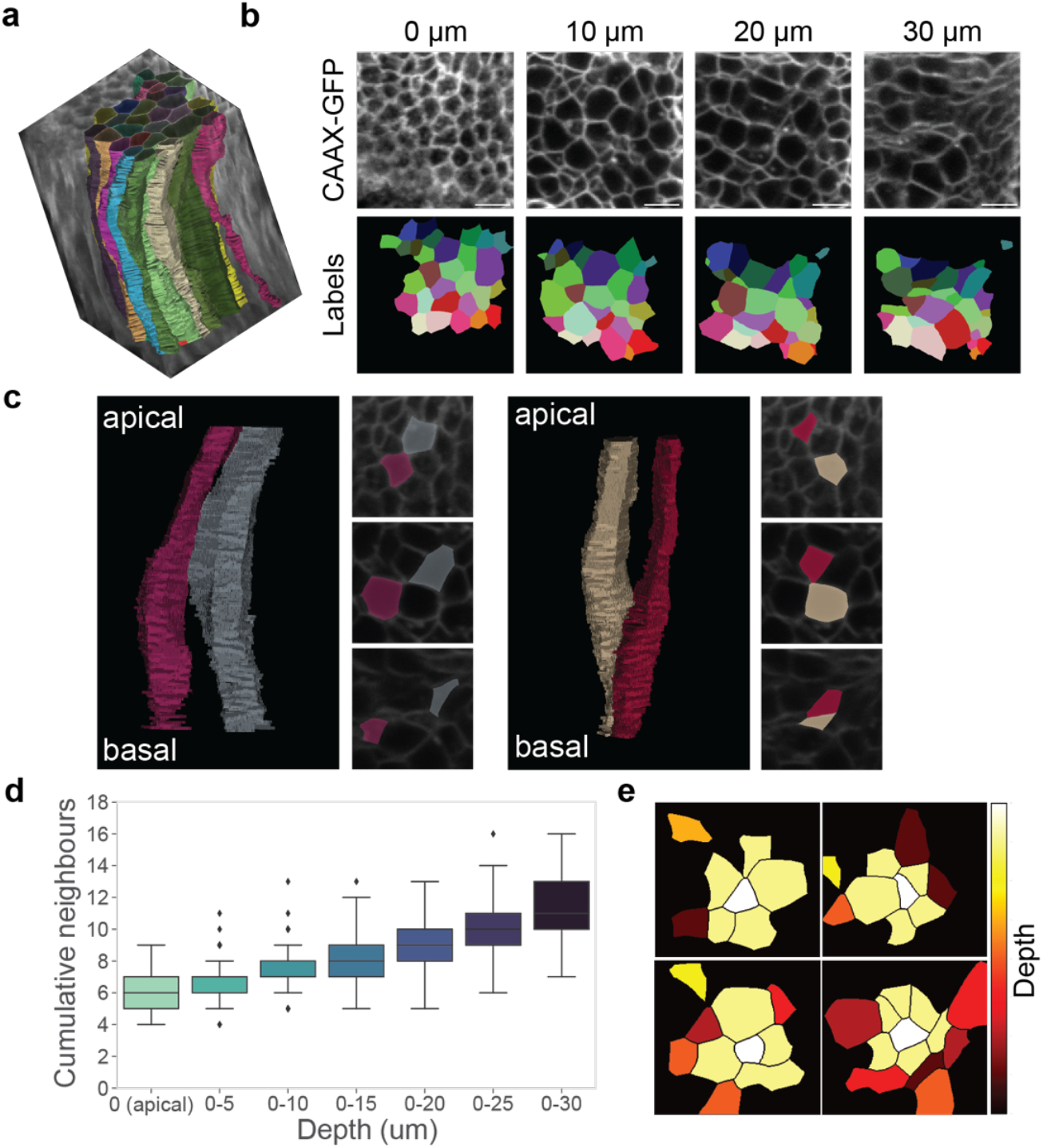
The topology of cell-cell contacts in 3D increases potential signalling range. **a**, Example 3D segmentation of wing disc cells. **b**, Membrane signal and corresponding segmented cell labels at different depths. Scale bar: 5 μm. **c**, Representative cell pairs showing apico-basal intercalations (apical neighbours moving further apart in z and vice-versa). **d**, Quantification of the number of cumulative cell neighbours along the cell depth (z), from N=3 wing discs. **e**, Apical labels of 4 representative cells (in white), showing their neighbourhood expansion in z. Neighbours of the cell in the middle are color-coded by the depth where they are encountered.

### A novel multi-layer signalling model explains dynamics of 3D signalling

To quantitatively assess how apical and lateral contacts shape Notch-Delta-mediated lateral inhibition, we extend Collier’s original framework (15) to a Multi-layer Signalling Model (MSM) that captures the full three-dimensional organisation of the epithelium. The model is constructed directly from 3D segmentations of wing imaginal discs, from which we extract the network of cell contacts at increasing depths, along with the shared edge lengths between neighbouring cells (Figure 3a). These contact interfaces are central to signalling, as the interaction strength between two cells scales with their contact boundary (20), providing a geometrically accurate representation of signalling potential.

**Figure 3:**
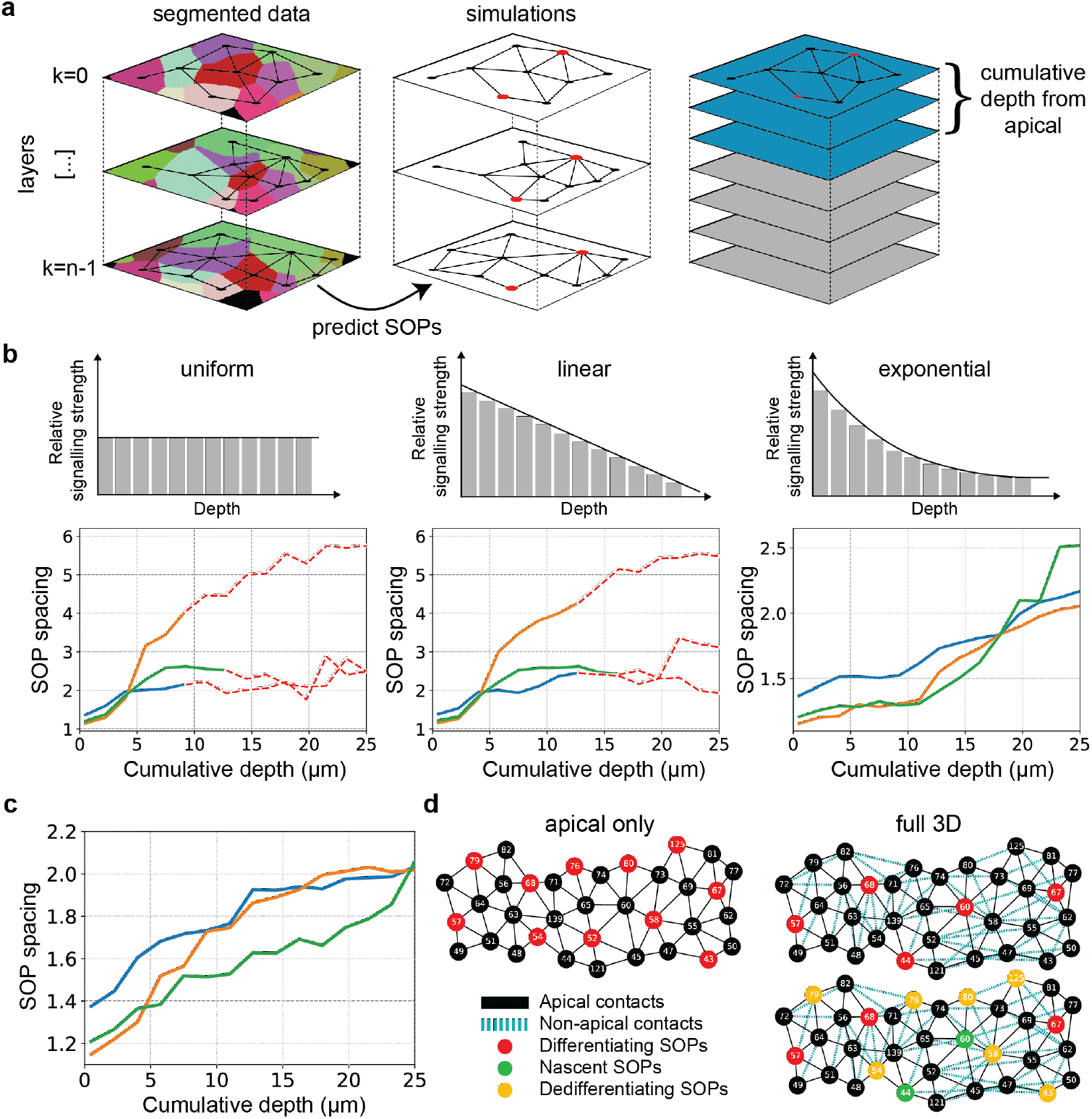
A multi-layer signalling model predicts the role of 3D contacts on SOP spacing. **a**, Generation of the Multi-layer Signalling Model. Left and middle panels: segmented data from experimental imaging is used to reconstruct the 3D cell contact network across *n* apico-basal layers. The MSM is then applied to predict SOP selection (red dots) by simulating Notch-Delta signalling across the whole 3D tissue architecture. Right panel: simulations are performed considering increasing cumulative depths in 3D to assess the role of lateral contacts. **b**, Top: Different simulated signalling profiles along the apico-basal axis. Bottom: SOP spacing as a function of cumulative signalling depth (simulations based on 3 different wing discs, corresponding to the three lines). Red dashed segments indicate degenerate patterning, including cases where SOP cells touch apically, patterning fails to emerge, or the number of SOPs is too low for a statistical analysis. Function parameters are listed in Supplementary Table 2. **c**, SOP spacing under signalling weights fitted to the experimentally measured Notch intensity profile (Supplementary Figure 1). **d**, Zoom-in of simulation predictions comparing apical-only (left) and full 3D signalling (right), with cells represented as nodes in a contact graph. Black edges denote apical contacts and turquoise dashed edges denote contacts that occur only in non-apical layers. Nodes in the bottom right graph are colour-coded to highlight cell fate changes compared to the apical only pattern.

The MSM captures the cumulative influence of neighbours located at different depths, with each layer contributing according to its relative signalling strength, reflecting Notch concentration along the apico-basal axis (Figure 1f). This approach enables us to adjust the balance between apical and deeper-layer inputs, revealing how 3D contacts collectively extend the range of lateral inhibition and modify SOP spacing. A detailed description of the model is provided in the Methods and Supplementary Information.

Firstly, we sought to systematically explore this effect by simulating three hypothetical depth-dependent signalling strength profiles: uniform, linear, and exponential (Figure 3b). In all cases, we ensured that apical-only signalling robustly produced a well-defined salt-and-pepper pattern, providing a consistent reference to assess how additional 3D contacts alter SOP spacing (see Methods and Supplementary Materials). Our results show that SOP spacing increases as the cumulative signalling depth grows for all profiles, but the rate and stability of this increase depend on how rapidly the signalling strength decays with depth. Uniform and linear profiles produce sparser patterns at greater depths but often result in degenerate outcomes: SOP cells touching apically, patterning failing to emerge, or the number of SOPs being too low for a statistical analysis. By contrast, the exponential profile yields a smoother and more biologically consistent increase in SOP spacing. We then quantified the distribution of Notch along the apico-basal axis in late third instar wing discs and found that it best fits an exponential distribution (Supplementary Figure 1). Fitting the experimental data yielded the final model (Figure 3c), which accurately captures the transition between salt-and-pepper (distance of 1) and sparse patterning (distance of 2) when considering increasing 3D layers.

Overall, we found that incorporating 3D contacts introduces additional inhibitory interactions that alter the fate of SOPs, refining the final pattern and increasing spacing (Figure 3d, top row). The change in SOP spacing arises because of dedifferentiation in crowded regions and the emergence of new SOPs in less inhibited areas (yellow and green nodes in Figure 3d).

### Altering 3D cell shapes affects SOP spacing

To further investigate the role of 3D contact topology on signalling, we performed *in silico* perturbation experiments to alter cell shapes by progressively “pruning” 3D contacts in the network and study the effect on SOP spacing (Figure 4a, see Methods for details). This approach simulates a “straightening” of the cells, reducing their tortuosity along the apico-basal axis. We found that, as the cells are progressively straightened, apico-basal intercalations are lost and the number of newly acquired neighbours in z decreases (Figure 4b, top). SOP spacing is initially not affected by mild straightening, but drops rapidly after 60% (Figure 4b, bottom).

**Figure 4:**
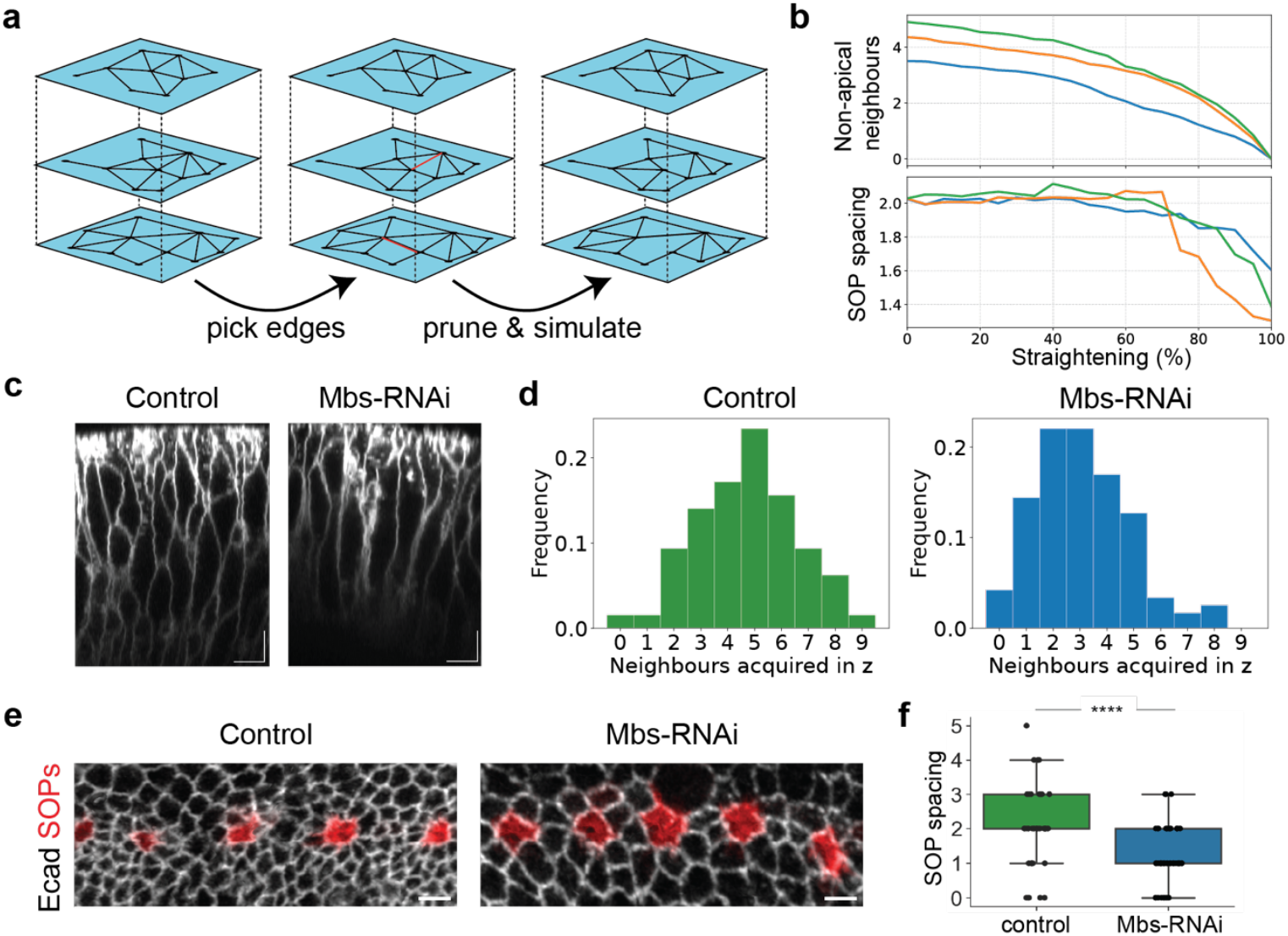
Changes in 3D contacts affect SOP patterning. **a**, Edge pruning approach to simulate the effect of altering 3D contacts, progressively “straightening” the cells. **b**, Effect of straightening the cells on the number of non-apical neighbours (top) and simulated SOP spacing (bottom). **c**, Side views of controls discs and Mbs-RNAi discs showing straighter lateral interfaces (scale bar, 5 μm). **d**, Quantification of neighbours acquired by cells in 3D in the two conditions, from N=3 wing discs each. **e**, SOP pattern images (scale bar, 5 μm) and **f**, SOP spacing quantification in the two conditions, from N=3 wing discs each. p-value<0.0001 from unpaired t-test.

To test the model predictions, we performed *in vivo* genetic modifications to directly perturb 3D cell shapes and quantified the impact on SOP spacing. We performed RNA interference (RNAi) against the myosin binding subunit (Mbs) of myosin phosphatase (which normally inactivates Myo II by dephosphorylating its regulatory light chain). Mbs-RNAi wing discs have previously been shown to display high cell edge tension and decreased tissue fluidity (fewer 2D intercalations over time) (21). We find that Mbs-RNAi wing discs have altered 3D shapes, with straighter lateral interfaces compared to control discs (Figure 4c). This altered cell shape also affects the topology of cell-cell contacts, with an overall decrease in the number of neighbours each cell contacts along the apico-basal axis (Figure 4d). While cells in control wing discs acquire 5 new neighbours through apico-basal intercalations on average, cells in Mbs-RNAi discs only contact 3 new cells on average, with some cells not changing neighbours at all.

To check if the altered 3D contacts induce a change in signalling and patterning, we imaged pre-pupal wing discs from Mbs-RNAi and control wing discs with the neuralised-RFP marker and anti-Ecad immunostaining to quantify the apical spacing of SOPs (Figure 4e). Focusing on the narrow pre-pupal developmental time window ensures that we are looking at the final SOP pattern. As predicted, we found that SOPs are more closely spaced in Mbs-RNAi discs compared to control discs (Figure 4f). Our results indicate that to fully capture the potential of cell-cell signalling, 3D cell topology must be taken into account.

## Discussion

Our work strongly highlights the need to consider 3D epithelial cell shapes in inter-cellular signalling: they control both the topology and the cell-cell interface areas across which contact signalling can act. Strikingly, we show that for apico-basally elongated cells, considering just the apical cell surface signalling strongly underestimates the full signalling potential of a cell. This is because the extensive lateral contacts can have a profound impact on signalling, even though a seemingly minor proportion of receptor/ligand is localised on them per unit area. Importantly, 3D cell topology is a readout of the history and mechanical status of a tissue. Thus, regulating intercellular signalling via 3D cell topology provides a novel mechanism of feedback control between mechanical and biochemical signalling (22).

In this study we have analysed “snapshots” of 3D cell-cell contacts, but how dynamic do we expect these to be? So far, it has not been possible to characterize dynamic apico-basal intercalations within live tissues over time; however, we speculate that lateral cell-cell contacts might be relatively dynamic. Interkinetic nuclear migration, where mitotic cells migrate apically to divide, is essential for successful mitosis in dense pseudo-stratified tissues (23) and will likely promote dynamic changes in 3D neighbours. In wing discs, however, we found that apico-basal intercalation extended the neighbourhood of cells to include second-row apical neighbours, but rarely beyond that (Figure 2e). Therefore, while we imagine that dynamic changes in 3D neighbours enable cells to “explore” their neighbourhood, this appears to occur without their effectively expanding the inhibition range. The fact that wing discs cells are tethered on both sides (basal attachments to the basement membrane and apical cell-cell junctions), together with fundamental limits to epithelial cell tortuosity, probably limits the maximum effective contact range that can be achieved. Having these multiple redundant but still spatially restricted contacts could indeed help confer robustness to signalling and the final developmental pattern, without the need for actively controlled processes like protrusions. Continuous improvements in imaging and segmentation may soon make it possible to address the question of contact dynamics experimentally and shed further light into the importance of 3D shape changes in signalling within tissues.

The novel mathematical modelling framework we have introduced here could readily be applied to other systems where contact topology and signalling are relevant. This is especially true now that widespread open-access AI tools like Cellpose (24) have made it easier to obtain 3D cell segmentations. The same framework can also be applied for systems where cell-cell contacts change over time in 2D due to cell movements, using a 2D timelapse instead of a 3D stack as input and with the option to “weigh” signalling over time as needed. Finally, it could be extended to include different types of contact-dependent or morphogen-based signalling.

The need to consider 3D cell topology in signalling has broad implications in development, patterning and growth control across multiple tissues and organisms. Cells with complex 3D shapes and apico-basal intercalations can be found across several organisms, especially in curved tissues. Examples include zebrafish (1) and sea star (25) embryos, *Drosophila* salivary glands (1), MDCK 3D cysts (26) and the mouse embryonic lung (2). This suggests that it could be a universal mechanism by which cell-cell signalling is tuned, especially within curved and pseudo-stratified tissues. Indeed, changing tissue height or cell shapes could be a way in which developing tissues gradually modify their effective cell-cell signalling distance to coordinate cell growth and fate specification (which are direct consequences of signalling) with cell and organ morphogenesis. Fine-tuning signalling range via regulating 3D cell topology may be a unified mechanochemical mechanism to ensure organs grow to the right size, shape and pattern.

## Extended Data

**Extended Data Figure 1:**
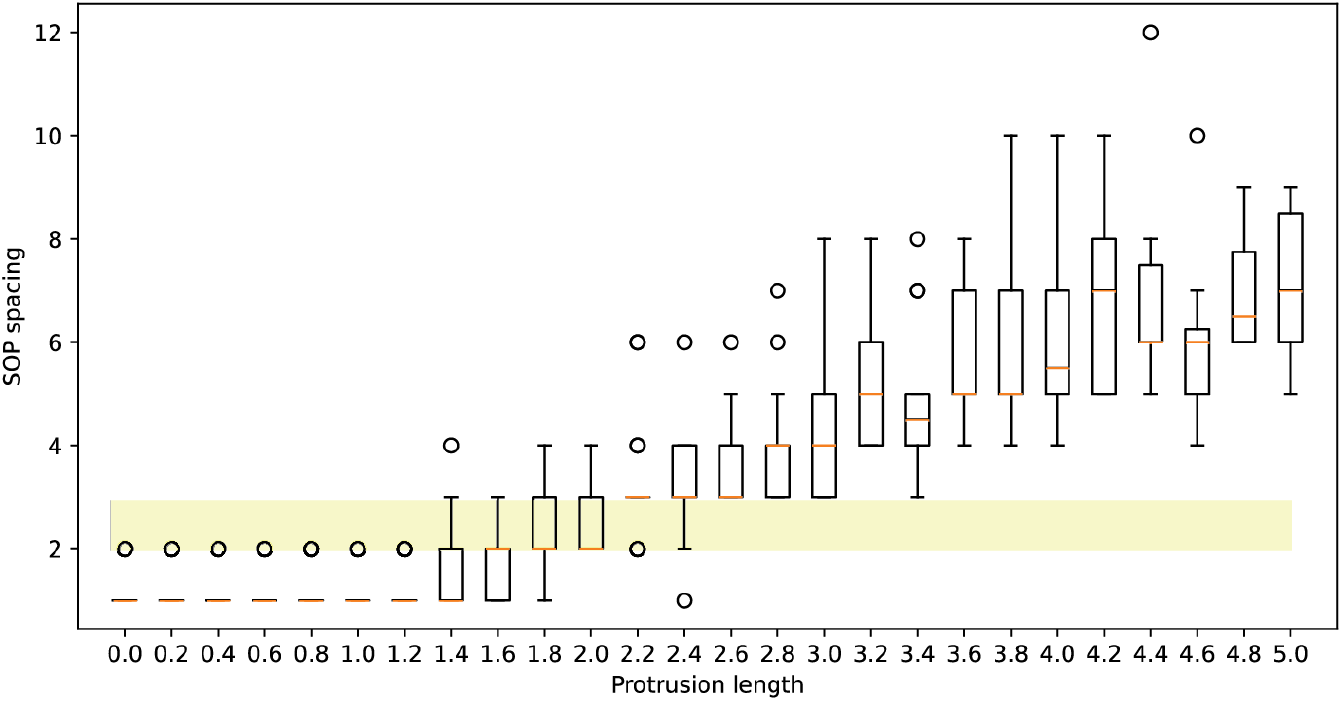
Long-range signalling range in 2D simulations. Results from 2D simulations of Notch-Delta signalling where the signalling range is tuned between 0 and 5 cell diameters (simulated as protrusion length). Note that to obtain an experimentally relevant range of SOP distances (2-3 SOP spacing, area shaded in yellow) the signalling range needs to be between 1.4 and 2.0. Here, we adapted the ϵ-Collier model of long-range signalling introduced by Berkemeier & Page (2023), with ϵ = 0.2.

**Extended Data Figure 2:**
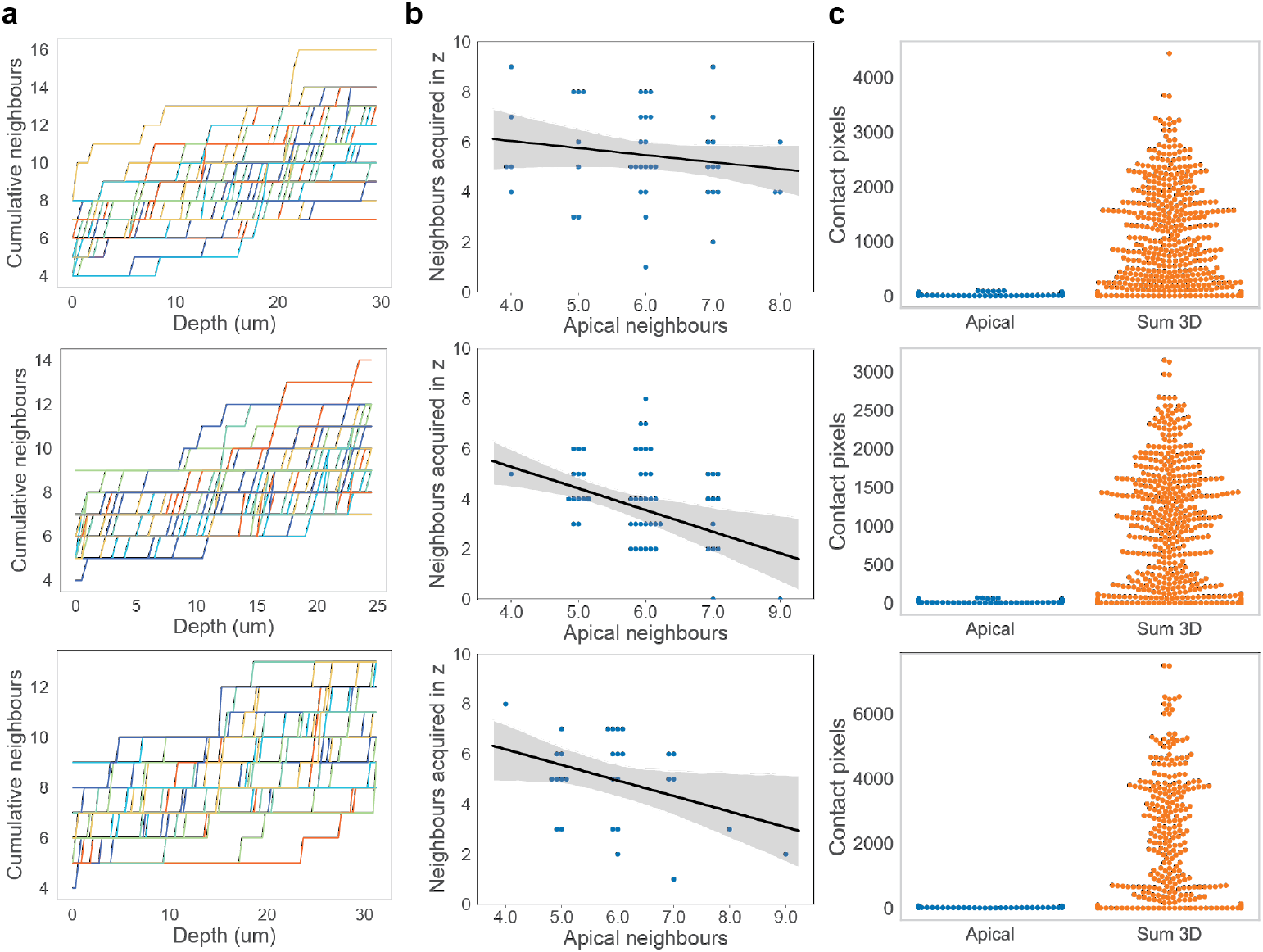
Relationship between apical and lateral cell-cell contacts. **a**, Increase of neighbours over depth: each line represents a cell, showing that all cells are scutoids and acquire new neighbours in z. **b**, Correlation between starting apical neighbours and additional neighbours gained in z, showing that cells that start with more apical neighbour gain slightly fewer neighbours in z. **c**, Comparison of cell-cell contact interfaces lengths apically and in 3D, showing that apical interfaces are overall much smaller due to the elongated shape of the cells. Each point corresponds to the interface between two cells. All data is shown for 3 different wing discs, each displayed on a separate row.

## Methods

### Fly husbandry

Flies were raised on a standard cornmeal molasses fly food medium at 25°C. Per 1L, the fly food contained 10g agar, 15g sucrose, 33g glucose, 35 g years, 15g maize meal, 10g wheat germ, 30g treacle, 7.22g soya flour, 1g nipagin, 5mL propionic acid.

For crosses, virgin females were placed in tubes with males at an approximate 2:1 ratio and flipped daily. Crosses were kept at 25°C.

### Fly stocks

The following fly stocks were used in this study: Ecad-GFP (27), neurPH-RFP (SOP membrane marker, gift of Schweisguth lab), Dl-GFP (28), Ni-GFP (17), yw; Ubi-GFP-CAAX (CAAX-GFP, DGRC 109824 FBID: FBtp0011013), NubGal4, UAS-myrGFP (nubbin-Gal4 FBID: FBti0016825 recombined with myrGFP, gift of Thompson group), UAS-Mbs-RNAi (II, VDRC KK Library) and Canton-S (Bloomington, 64349).

### Dissections, wing disc ex-vivo culture and sample mounting

Male and female larvae were dissected at late 3^rd^ instar development (approximately 110-120hr AEL) for experiments apart from endpoint pattern spacing quantification of control vs Mbs-RNAi vs control discs, for which pre-pupae were used. Larvae were rinsed in PBS and dissected in culture media: Shields and Sang M3 media (Merck) supplemented with 2% FBS (Merck), 1% pen/strep (Gibco), 3 ng/ml ecdysone (Merck) and 2 ng/ml insulin (Merck).

For fixed sample imaging, wing discs were mounted on a standard coverslip using Fluoromount G slide mounting medium (SouthernBiotech) between two strips of double-sided tape as a spacer. For live imaging (upright 2-photon microscope), wing discs were mounted on a coverslip between two strips of double-sided tape as a spacer and filled with culture media. Alternatively, for imaging on an inverted confocal microscope, discs were mounted in a 35mm glass-bottom dish filled with culture media by creating a small round well with double-sided tape and sealing the top with filter paper.

### Immunohistochemistry and tissue staining

Discs were dissected in culture media and fixed for 30 min in 4% paraformaldehyde, washed 4×10 min with PBT (PBS, 0.3% Triton X-100), blocked for 1 h with 0.5% BSA PBT and stained with primary (overnight at 4°C), washes were repeated before incubating in fluorescently conjugated secondary antibodies (1h at room temperature) and were washed 3×20 min PBT, followed by 3x quick washes in PBS.

Antibodies used: primary rat anti-E-cad (Developmental Studies Hybridoma Bank) (1:50 dilution) and secondary anti-rat Cy5 (1:400 dilution).

Staining with Cell Mask DeepRed was performed live by incubating discs in a small volume (20 μl) containing 0.5 μl undiluted stock solution in culture media for 20 min at room temperature.

### Sample Imaging (confocal)

For initial SOP pattern quantification (Figure 1d), fixed wing discs were imaged on a spinning disc confocal (Zeiss AxioObserver Z1 inverted stand attached to a Yokogawa CSU-W1 spinning disc scan head) using a 63x oil objective and Prime 95B Scientific CMOS camera. Z-stack were acquired with a step of 0.5 μm to capture the apical domain of the wing discs. For live imaging of Notch/Delta localisation, sample were imaged on a Zeiss LSM900 confocal microscope with a 63x oil (NA=1.4) objective. A large field of view was acquired to image most of the wing disc pouch using a zoom of 1, laser power of 1% and Airyscan detector. Z-stack were acquired with a step of 0.5 μm up to a depth of around 40 μm.

For SOP spacing comparison between control and Mbs-RNAi wing discs, fixed samples were imaged on a Zeiss LSM900 confocal microscope with a 63x oil (NA=1.4) objective. Field of views were adjusted to image most of the SOP pattern using a zoom of 0.8, laser power of 1% and Airyscan detector. Z-stack were acquired with a step of 0.5 μm up to a depth of around 40 μm.

### Sample Imaging (two-photon)

Two-photon images were taken on a LEICA DIVE confocal microscope with a 63x oil objective, zoom of 8 using a wavelength of 924 nm for two-photon GFP excitation. Z-stack were acquired with a step of 0.3 or 0.5 μm to capture as deep as possible in the tissue until excessive signal degradation (typically 30 μm). Z-compensation on both the Excitation and Emission Gains was applied to compensate for the loss of intensity at depth within the tissue.

### SOP spacing quantification from experiments

The adaptive projection in EpiTools (29) was used to selectively project the apical surface of wing discs. Ecad was used as the reference channel to identify the apical plane and selectively project the apical neuralised signal. The projected Ecad image was segmented in EpiTools and SOPs were manually picked based on the neuralised signal intensity. SOP spacing (number of cells between each SOP) was computed automatically from the segmentation as the shortest path between identified SOPs using a python script. Prepupal discs were occasionally too curved to obtain a full correct projection, in which case only the analysis was restricted to the correctly projected regions.

### Quantification of apical properties

The projected and segmented images from the previous step were processed in napari to extract measurement tables using the regionprops plugin. Quantification of cell properties from the exported tables was done with python scripts.

### 3D segmentation and analysis

Images were segmented using Cellpose (24) (model:cyto 3, parameters: cell diameter = 60 (70 for Mbs-RNAi, flow threshold = 0.4, cell probability threshold = 0 and stitch threshold = 0.4). Manual corrections were done in napari (30), see (19) for more details. Quantification of cell properties (neighbours, size) was done using python scripts after exporting cell properties in napari using the EpiTools (31) and regionprops plugins.

### Quantification of apico-basal Notch distribution

A square ROI was selected in Fiji to analyze the central part of the wing pouch (708 x 708 pixels). A mask was created from the cell membrane staining (cellMask) to restrict Notch quantification only to the membrane, excluding intracellular Notch: briefly, images were processed in Fiji by applying a Gaussian Blur (radius of 1), Thresholding using Otsu’s method and dividing the resulting image by 255 to obtain a mask. The mask image was then multiplied to the Notch channel using the Image Multiplication operation. A correction factor was calculated from the cellMask signal to correct for signal loss in z due to optical attenuation, and then multiplied to the Notch signal. The final scaled and masked Notch intensity was computed from three different wing discs and fitted to an exponential function (Supplementary Figure 1a). For Fig. 1f panels, individual slices showing the lateral view of cells were background-subtracted (rolling ball in Fiji with a radius of 50) to improve clarity.

### Multi-layer Signalling Model

The dynamics of Notch-Delta signalling within each cell *i* = 1, …, *N* may be represented by the following system of equations (15):

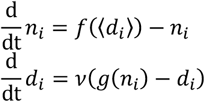

We take *f, g*: [0, ∞) → [0, ∞) to be continuous increasing and decreasing Hill functions, respectively,

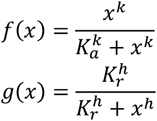

for *x ≥* 0, where *k* and *h* (Hill coefficients) determine the steepness of the response, and *K*_*a*_ and *K*_*r*_ set the half-maximal activation and repression thresholds, respectively. *ν >* 0 is the ratio between Notch and Delta decay rates. To simulate three-dimensional interactions, we introduce

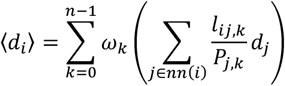

for a total number of signalling layers *n* (layer range), where, for each layer *k* = 0, …, *n* − 1, *l*_*ij*,*k*_ is the length of the shared edge between cells *i* and neighbouring cell *j*, and *P*_*j*,*k*_ *is* the cross-sectional perimeter of cell *j* at that layer. *nn*(*i*) is the set of nearest neighbours of cell *i*, and *ω*_*k*_ is the signalling weight of layer *k*. The total number of signalling layers can be defined by *n = L* / Δ*L*, where *L* is the actual apical-to-basal length, determined experimentally, and Δ*L* is the width of each layer. To guarantee robust patterning and avoid degenerate regimes, we set *k* = 2, *h* = 8, *K*_*a*_ = 10^−1^, *K*_*r*_ *=* 10^−3^, and *ν* = 0.09.

The signalling weights {*ω*_*k*_} were derived from masked Notch intensity profiles measured along the apico-basal axis (Δ*L =* 0.5 μm, total depth *L* = 32 μm). A non-increasing function ω(*z*) was fitted to the intensity data, and weights for each layer *k* were computed as

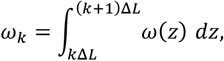

ensuring that the total weight matches the integrated signalling intensity (Supplementary Figure 1). We consider alternative functional forms of *ω*(*z*) (Figures 3b,c), selected so that *ω*_0_ yields a minimum value required for robust pattern formation.

Cells are considered SOPs if their Delta activity exceeds a threshold of 0.8. To quantify SOP spacing, we construct an unweighted graph from the apical centroids, where nodes represent cells and edges denote apical contacts. The shortest path between SOPs in this graph gives the number of intervening non-SOP cells, providing a spacing measure that reflects actual contact geometry. To study variation along the anterior-posterior axis, we divide the apical layer into horizontal bands of fixed dorso-ventral height, chosen so that each band contains a comparable number of apical centroids across discs. The mean SOP spacing is computed within each band and combined into a moving-average profile, which reveals that increasing the number of signalling layers *n* systematically enlarges SOP separation (Figure 3; Supplementary Figure 2).

To assess how straightening of the epithelium influences SOP patterning, and to enable a direct comparison with the Mbs-RNAi experiments (Figure 4), we introduce a straightening model that progressively reduces the contribution of non-apical contacts. This is achieved by interpolating each layer’s adjacency toward the apical contact map, controlled by a parameter α ∈ [0,1]. At α = 0, the full three-dimensional connectivity is preserved, whereas at α = 1 all layers are forced to match the apical configuration, resulting in fully “straight” cells. For intermediate values, edges present only in deeper layers are pruned and missing apical edges are restored, with the extent of these edits determined by the displacement of cell centroids relative to their apical positions. This approach allows us to systematically quantify the role of non-apical neighbours in lateral inhibition and SOP spacing, bridging the gap between raw 3D structure and a fully straight model. The detailed mathematical formulation is provided in the Supplementary Information.

Each SOP spacing plot shows the mean of 500 simulations, computed for every layer depth (Figures 3b,d) and straightening factor α (Figure 4b, as a percentage).

## Data Availability

3D images and segmented labels will be uploaded on the BioImage Archive https://www.ebi.ac.uk/bioimage-archive/. Other data is available upon request.

## Code Availability

Code is available at the repository: https://github.com/fberkemeier/MultiLayer-NotchDelta.

## Acknowledgements

G.P. was supported by an EMBO Long-Term Fellowship (ALTF 786-2020). F.B. was supported by a UCL PhD Studentship Award from the Department of Mathematics, the Leverhulme Trust Research Project Grant RPG-2022-028, and a Rokos Postdoctoral Associate position at Queens’ College, Cambridge. Y.M. was supported by the MRC award MR/W027437/1, a Lister Institute Research Prize and EMBO Young Investigator Programme. B.B. receives support from the Medical Research Council - Laboratory of Molecular Biology (MC_UP_1201/27). We thank R. Barrientos, P. Vicente Munuera, Profs. R. Pérez-Carrasco, P. Pearce and M. Dalwadi for their constructive and insightful feedback, which greatly improved the manuscript.

## Author contributions

Y.M. and G.P. conceived the project. G.P. designed, performed and analysed experiments. F.B. designed and developed the model. G.P., F.B., B.B, K.M.P. and Y.M. all contributed to experimental design, writing the manuscript and preparing the figures.

