## Supplementary Information for "3D epithelial cell topology tunes signalling range to promote precise patterning"

### 1 A multi-layer model of Notch-Delta signalling

The dynamics of Notch-Delta signalling within each cell $i$ may be represented by the following system (Collier et al., 1996)

$$\frac{d}{dt}n_i = f(\langle d_i \rangle) - n_i \tag{1}$$

$$\frac{d}{dt}d_i = \nu(g(n_i) - d_i) \tag{2}$$

for $1 \leq i \leq N$, where $N$ is the total number of cells. We define $f$ and $g$ as Hill functions

$$f(x) = \frac{x^k}{K_a^k + x^k}, \quad g(x) = \frac{K_r^h}{K_r^h + x^h} \tag{3}$$

where $k$ and $h$ (Hill coefficients) determine the steepness of the response (cooperativity), and $K_a$ and $K_r$ set the half-maximal activation and repression thresholds, respectively. $\nu > 0$ is the ratio between Notch and Delta decay rates. To simulate three-dimensional interactions, we introduce

$$\langle d_i \rangle = \sum_{k=0}^{n-1} \omega_k \left( \sum_{j \in \mathbf{nn}(i)} \frac{\ell_{ij,k}}{P_{j,k}} d_j \right) \tag{4}$$

for a total number of signalling layers $n$ (layer range), where, at each layer $k$ ($0 \leq k \leq n-1$), $\ell_{ij,k}$ is the length of the shared edge between cells $i$ and neighbouring cell $j$, and $P_{j,k}$ is the cross-sectional perimeter of cell $j$ at that layer. $\mathbf{nn}(i)$ is the set of nearest neighbours of cell $i$, and $\omega_k$ is the signalling weight of layer $k$. The total number of signalling layers can be defined by $n = L/\Delta L$, where $L$ is the actual apical-to-basal length, determined experimentally, and $\Delta L$ is the width of each layer.

### 2 Model parameters

To ensure we sit in a non-degenerate regime that avoids both uniform inhibition and runaway oscillations, and to strike a balance between apical versus deeper-layer signalling, we carried out a numerical sensitivity sweep over $(k, h, K_a, K_r, \nu)$. We then selected

$$k = 2, \ h = 8, \ K_a = 10^{-1}, \ K_r = 10^{-3}, \ \nu = 0.09 \tag{5}$$

as lying robustly within the region where clear, stable salt-and-pepper patterns form and where neither apical nor lateral signals alone dominate.

Regarding the signalling weights $\{\omega_k\}$, we rely on the masked Notch intensity measured along the cells' apico-basal axis, with a layer resolution of $\Delta L = 0.5$ µm, spanning $L = 32$ µm. We fit a non-increasing signalling weight function $\omega(z)$ to the measured intensity values (Supplementary Figure 1a). We bin the signal intensity across the different wing discs, defined by their respective layer ranges (Supplementary Figure 1b). For each disc, we determine the bins by setting their width equal to the layers' height, $\Delta L$. Given the total number of layers in each disc, $n$, this setup spans a vertical depth of $n\Delta L$. Then, within each bin, we define the signalling weights $\omega_k$ as the area under the fitted curve $\omega(z)$, that is,

$$\omega_k = \int_{k\Delta L}^{(k+1)\Delta L} \omega(z)\, dz, \tag{6}$$

ensuring that the total weight across all layers is consistent with the integrated signalling intensity. Note that the total signalling weight, i.e., the integral of the depth-dependent profile $\omega(z)$ over all $n$ layers, can be re-scaled simply by adjusting the activation threshold $K_a$ in Eq. (3). Increasing $K_a$ reduces $f$ and thus uniformly lowers the effective total weight without altering the shape of $\omega(z)$. This approach provides a robust and consistent measure of Notch signalling strength, capturing the signal distribution in a manner that respects the specific geometry and size of each wing disc.

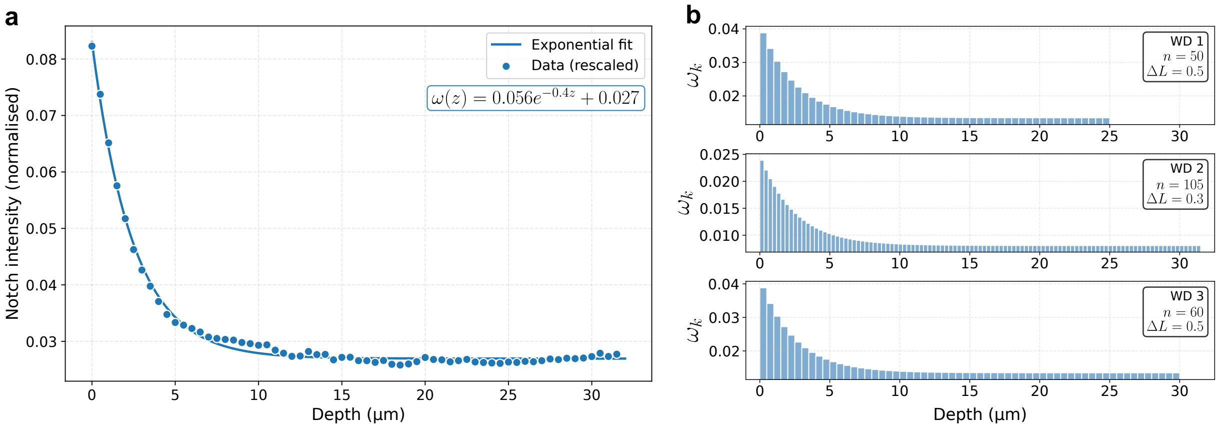


**Supplementary Figure 1.** **a** Masked Notch intensity data is fitted to a normalised curve $\omega(z) = Ae^{-Bz} + C$, with parameters $(A, B, C) \approx (0.056, 0.4, 0.027)$. This dataset comprises 64 data points, spanning a total depth of $0.5 \times 64 = 32$ µm. **b** Distributions of signalling weights for three different wing discs, each with varying total number of frames ($n$) and layer widths ($\Delta L$), calculated using Eq. (6).

### 3 SOP cell spacing

To quantify apical SOP spacing, we use the apical centroids of all cells to construct an unweighted graph where nodes represent cells and edges denote apical contacts between neighbouring cells. For each SOP cell, we identify its closest SOP neighbour by computing the shortest path in this graph (Supplementary Figure 2a). The resulting path length corresponds to the number of apical cell-to-cell steps between SOPs, effectively counting the non-SOP cells that separate them. This graph-based measure captures the spacing of SOPs in terms of actual contact geometry rather than simple Euclidean distances. In each simulation, SOP cells are identified as those with Delta activity exceeding 0.8. To study how SOP spacing varies along the antero-posterior direction, we sample the apical layer with rectangular bands of fixed dorso-ventral height. This choice is motivated by the observation that SOP cells tend to align approximately along the antero-posterior axis, so averaging within narrow dorso-ventral strips preserves this directional structure. Band heights are chosen so that, on average across all wing discs, each band contains the same number of apical centroids which limits sampling bias. Within each band, we compute the mean nearest neighbour distance between SOPs whose apical centroids fall inside it, and by sliding the band along the antero-posterior axis we obtain a moving average profile of SOP spacing (Supplementary Figure 2b). This procedure smooths local fluctuations and reveals that increasing the number of signalling layers $n$ systematically enlarges the average SOP separation, consistent with the role of three dimensional contacts in setting the pattern (Figure 3).

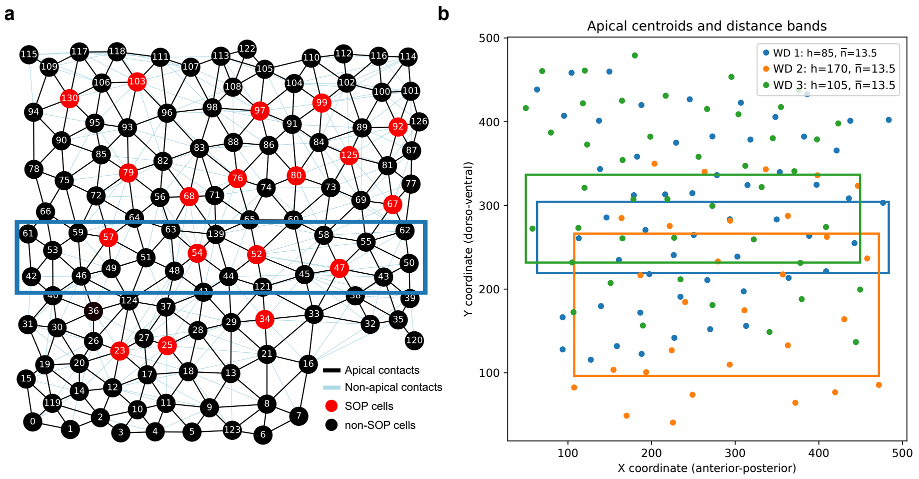


**Supplementary Figure 2.** **a** Example of a simulated pattern where SOP cells (in red) are represented as nodes (centroids) of a multi-layered graph. Black edges indicate apical contacts, while light blue edges indicate exclusively non-apical contacts. SOP spacing is calculated by measuring the shortest-path distance between SOP nodes, averaged within sliding bands (blue rectangle) along the dorso-ventral (vertical) axis of the tissue. **b** Apical centroids of SOP cells and horizontal (antero-posterior) distance bands used to compute graph-based SOP spacing. Each colour corresponds to a different wing disc, with the mean number of apical centroids per band indicated ($\bar{n} = 13.5$). Bands are defined along the dorso-ventral axis so that each contains, on average across all wing discs, the same number of apical centroids, ensuring consistent sampling for the moving average of SOP spacing. Heights were set, respectively, to $(h_1, h_2, h_3) = (85, 170, 105)$.

### 4 Straightening model

Wing imaginal discs are composed of highly elongated epithelial cells whose apico-basal axes can become tortuous in deeper layers. As a result, their lateral contacts in depth often deviate from the packing observed at the apical surface, leading to enhanced lateral inhibition and sparser SOP cell patterning, as discussed in the main text.

To further understand the effect of tortuosity in cell differentiation, we introduce a straightening model that allows us to interpolate between each layer's raw, three-dimensional contacts and the apical adjacency by tuning a single parameter $\alpha \in [0, 1]$. When $\alpha = 0$, every layer remains exactly as measured; when $\alpha = 1$, every layer is forced to match the apical contact map; intermediate values gradually prune spurious contacts and restore missing ones based on each cell's deviation from the apical plane.

First, we convert each (edge-weighted) adjacency matrix $A^{(k)} \in \mathbb{R}^{N \times N}$ into a binary contact map, for each layer $k = 0, 1, \ldots, n-1$,

$$b_{ij}^{(k)} = \begin{cases} 1 & \text{if } A_{ij}^{(k)} > 0 \\ 0 & \text{otherwise} \end{cases} \tag{7}$$

so that $b_{ij}^{(k)} = 1$ if and only if cells $i$ and $j$ touch in layer $k$. We also record each cell's two-dimensional centroid $\mathbf{p}_i^{(k)} \in \mathbb{R}^2$ and compute its radial displacement from the apical layer

$$d_i^{(k)} = \left\|\mathbf{p}_i^{(k)} - \mathbf{p}_i^{(0)}\right\|, \tag{8}$$

which measures how far cell $i$ has deviated from the apical surface.

Next we identify two types of discrepancies between layer $k$ and the apical layer $k = 0$. *Extraneous* edges are those contacts present only in depth but absent apically,

$$E_{\text{ex}}^{(k)} = \{(i, j) \mid b_{ij}^{(0)} = 0,\ b_{ij}^{(k)} = 1\}, \tag{9}$$

and *missing* edges are those contacts present apically but lost in depth,

$$E_{\text{mi}}^{(k)} = \{(i, j) \mid b_{ij}^{(0)} = 1,\ b_{ij}^{(k)} = 0\}. \tag{10}$$

To each candidate $(i, j)$ we assign a severity score

$$\delta_{ij}^{(k)} = \max\big(d_i^{(k)},\ d_j^{(k)}\big), \tag{11}$$

so that contacts between cells with greater deviation from their apical positions carry higher $\delta$.

We then set two quantile-based thresholds. From the set of extraneous scores at layer $k$ we take the $(1-\alpha)$-quantile

$$\tau_{\text{ex}}^{(k)}(\alpha) = Q_{1-\alpha}\big(\{\delta_{ij}^{(k)} : (i, j) \in E_{\text{ex}}^{(k)}\}\big), \tag{12}$$

and from the missing-edge scores at layer $k$ we take the $\alpha$-quantile

$$\tau_{\text{mi}}^{(k)}(\alpha) = Q_{\alpha}\big(\{\delta_{ij}^{(k)} : (i, j) \in E_{\text{mi}}^{(k)}\}\big). \tag{13}$$

This way, we are able to remove exactly an $\alpha$-fraction of the worst (highest-severity) extraneous edges, and add back exactly an $\alpha$-fraction of the best (lowest-severity) missing edges. If a category is empty, its threshold is set to $+\infty$ or $-\infty$ so that no edges are edited.

Finally, the straightened adjacency $\widetilde{A}_{ij}^{(k)}(\alpha)$ is given by

$$\widetilde{A}_{ij}^{(k)}(\alpha) = \begin{cases} 0, & \text{if } b_{ij}^{(0)} = 0,\ b_{ij}^{(k)} = 1,\ \delta_{ij}^{(k)} > \tau_{\text{ex}}^{(k)}(\alpha), \\ A_{ij}^{(0)}, & \text{if } b_{ij}^{(0)} = 1,\ b_{ij}^{(k)} = 0,\ \delta_{ij}^{(k)} < \tau_{\text{mi}}^{(k)}(\alpha), \\ A_{ij}^{(k)}, & \text{otherwise,} \end{cases} \tag{14}$$

so that at $\alpha = 0$ nothing changes, and at $\alpha = 1$ all extraneous edges are pruned and all apical edges restored, smoothly morphing each layer's measured connectivity toward the apical packing, and gradually removing non-apical neighbours (Figure 4; Supplementary Figure 3).

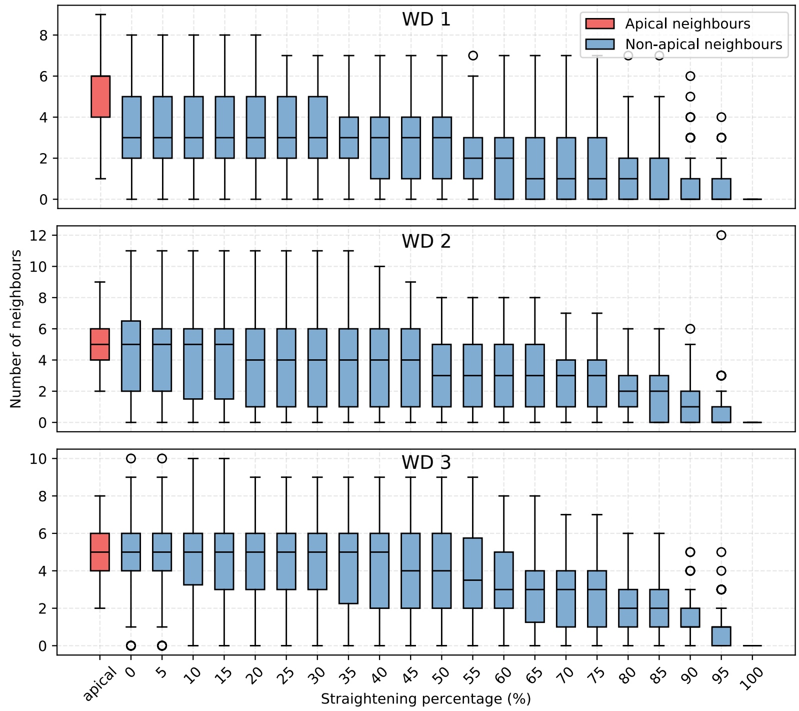


**Supplementary Figure 3.** Number of neighbours as a function of the straightening factor $\alpha$ (as a percentage).

### 5 Alternative models of long-range signalling

Abstracting away the full 3D structure of epithelial cells, our multi-layer model can be viewed as a mechanism for effective long-range signalling projected onto the apical surface. In such a 2D abstraction, extended lateral inhibition may also be achieved through protrusions, such as basal actin-based filopodia, which connect non-neighbouring cells. While wing disc cells have no visible baso-lateral filopodia, protrusion-mediated signalling has been shown to influence sparse SOP patterning in various contexts, particularly in the fly notum.

Several mathematical models have been proposed to describe such protrusion-driven interactions. Notably, Berkemeier and Page (2023) developed a framework for coupling juxtacrine and long-range signalling on a hexagonal lattice, introducing the $\epsilon$-Collier model. This model extends Collier's classical lateral inhibition scheme by incorporating a weighting factor, $\epsilon \in [0, 1]$, which balances local (juxtacrine) and non-local (protrusion-based) signalling contributions.

Mathematically, the main difference from the MSM is in the way we define the coupling term $\langle d_i \rangle$. In the $\epsilon$-Collier model, the coupling term typically separates into *short-range* and *long-range* components

$$\langle d_i \rangle = (1 - \epsilon) \sum_{j \in \mathbf{nn}(i)} \omega_J(i, j) d_j + \epsilon \sum_{j \in \mathbf{np}(i)} \omega_P(i, j) d_j, \qquad (15)$$

where $\mathbf{nn}(i)$ is the set of immediate neighbours and $\mathbf{np}(i)$ denotes the set of cells connected to $i$ via protrusions. $\omega_J(i, j)$ and $\omega_P(i, j)$ are non-negative weighting functions that specify how strongly cell $i$ samples Delta from cell $j$ through, respectively, direct (juxtacrine) contact and protrusions, with $\omega_P$ also a function of the protrusion length. They are typically normalised to reflect signalling/ligand availability.

When $\epsilon = 0$, the original short-range Collier model is recovered; increasing $\epsilon$ introduces progressively stronger long-range interactions that promote sparser, larger-scale patterns, as seen in Figure 1d and Extended Data Figure 1 of the main text. While our 3D contact-based MSM focuses on the geometry and depth of real cellular interfaces, the protrusion-based approach offers a complementary, more abstract view of long-range signalling. For further mathematical details and stability analysis of the $\epsilon$-Collier model, see Berkemeier and Page (2023).

### 6 Supplementary tables

The following tables summarise the key parameters used in our analysis. Table 1 lists the data-based structural properties of the wing discs, including the number of signalling layers, layer height, total depth, and the number of signalling cells. Table 2 reports the parameters of the signalling weight profiles $\omega(z)$ used across different models, highlighting the values of $A$, $B$, and $C$ for each functional form (Figure 3).

| **Wing disc** | **Layer range** ($n$) | **Layer height** ($\Delta L$) | **Depth** ($L$) | **Signalling cells** ($N$) |
|---|---|---|---|---|
| WD 1 (14/03) | 50 | 0.5 µm | 25 µm | 55 |
| WD 2 (17/03) | 105 | 0.3 µm | 31.5 µm | 29 |
| WD 3 (21/03) | 60 | 0.5 µm | 30 µm | 44 |

**Supplementary Table 1.** Wing disc data parameters.

| | **Uniform** | **Linear** | **Exponential** | **Optimal** |
|---|---|---|---|---|
| $\omega(z)$ | $C$ | $Az + C$ | $Ae^{-Bz} + C$ | $Ae^{-Bz} + C$ |
| Parameters | $C = 36$ | $A = -1.14$<br>$C = 36$ | $A = 10$<br>$B = 0.2$<br>$C = 2$ | $A = 40$<br>$B = 0.4$<br>$C = 0$ |

**Supplementary Table 2.** Signalling weight parameters across different profiles of $\omega(z)$. In all models, we set $L = 32$, and distance heights to $(h_1, h_2, h_3) = (85, 170, 105)$.
